# Rapid eye and hand responses in an interception task are differentially modulated by context-dependent predictability

**DOI:** 10.1101/2024.07.11.603058

**Authors:** Jolande Fooken, Parsa Balalaie, Kayne Park, J. Randall Flanagan, Stephen H. Scott

## Abstract

Humans can quickly generate eye and hand responses to unpredictable changes in the environment. Here, we investigated eye-hand coordination in a rapid interception task where human participants used a virtual paddle to intercept a moving target. The target moved vertically down a computer screen and could suddenly ‘jump’ to the left or right. In high-certainty blocks, the target always jumped, and in low-certainty blocks, the target only jumped in a portion of trials. Further, we manipulated response urgency by varying the time of target jumps, with early jumps requiring less urgent and late jumps requiring more urgent responses. Our results highlighted differential effects of certainty and urgency on eye-hand coordination. Participants initiated both eye and hand responses earlier for high-certainty compared to low-certainty blocks. Hand reaction times decreased and response vigor increased with increasing urgency levels. However, eye reaction times were lowest for medium-urgency levels and eye vigor was unaffected by urgency. Across all trials, we found a weak positive correlation between eye and hand responses. Taken together, these results suggest that the limb and oculomotor systems use similar early sensorimotor processing; however, rapid responses are modulated differentially to attain system-specific sensorimotor goals.

## Introduction

The ability to interact with moving objects in dynamic and unpredictable environments requires complex coordination between the eyes and hands. In a game of table tennis, players track the rapidly moving ball with their eyes and predict the ball’s trajectory after it hits the table to successfully intercept it. In such dynamic tasks, eye and hand movement control is shaped by how certain the available sensory information is and how urgent the motor response has to be executed. For example, there can be uncertainty in the direction the ball will travel following the initial bounce on the table (i.e., if the opponent puts a spin on the ball). There can also be variability in urgency as the opponent can hit the ball further or nearer to the player requiring less or more urgent responses, respectively. Thus, goal-directed responses often require rapid updating of visual and proprioceptive feedback to ensure successful motor actions.

In everyday action tasks, eye and hand movements have been shown to be coordinated in stereotypical ways (Land, 2006). In stationary reaching, eye movements typically lead hand movements, fixating on key landmarks and objects before they are manipulated (de Brouwer et al., 2021; Johansson et al., 2001). This relationship is more complex when interacting with moving targets. Intercepting a moving target involves predicting where the target will be at future states (Bosco et al., 2015; Fooken et al., 2021; Zago et al., 2010). This prediction is crucial to overcome the delays in our sensorimotor system (Brenner & Smeets, 2018; Todorov & Jordan, 2002; Zago et al., 2009). During interception tasks, maintaining gaze on the moving target (via smooth pursuit eye movements) allows continuous updating of the target’s trajectory that can aid motion prediction (Spering et al., 2011) and manual interception performance (Fooken et al., 2016; Goettker et al., 2019; Kreyenmeier et al., 2022; Mrotek, 2013; Mrotek & Soechting, 2007). However, eye movement behaviour is also affected by the predictability of the moving object’s path and visual certainty. In situations of high uncertainty, such as when the moving target bounces or is occluded, observers shift their gaze away from the moving object to the anticipated interception locations (de la Malla et al., 2019; Diaz et al., 2013; Fooken & Spering, 2019; Mann et al., 2019). Thus, the coordination between eye and hand movements can be modulated by visuomotor task demands.

Whereas a close coordination between eye and hand movements has been observed in action tasks, such as object manipulation or manual interception, the link between eye and hand movement control in tasks requiring rapid visuomotor responses is less clear. Previous research has shown that humans are able to elicit very rapid motor responses to sudden visual stimuli —a phenomenon referred to as express visuomotor responses or stimulus locked responses (Corneil et al., 2008). Express visuomotor responses have been found in the generation of saccadic eye movements (Dorris et al., 1997) as well as in the upper limb responses following visual target appearance (Pruszynski et al., 2010). For upper limb responses, it has been shown that high certainty about a forthcoming perturbation and temporal predictability of perturbation onset evokes reliable and rapid muscle responses (Contemori et al., 2021; Kozak et al., 2020). Early waves of upper limb muscle responses are also modulated by the level of response urgency, with movement corrections occurring earlier relative to the moment of perturbation when the time available to make a response is limited (Crévécoeur et al., 2013; Maurus et al., 2023; Poscente et al., 2021). The response modulation by both jump certainty and urgency suggests an influence of top-down control on express visuomotor responses, despite previous work highlighting their reactive nature (Corneil & Munoz, 2014).

Here, we examined eye and hand responses to visual perturbations during a rapid interception task. Participants used a virtual paddle to intercept a target moving down a vertically-oriented computer screen. In the majority of trials, the moving target was visually perturbed (‘jumped’), suddenly shifting spatial location to the left or right and continuing moving down after the jump. To examine the effect of jump certainty, we modified the frequency at which the target jumped. In the high-certainty condition, the target jumped to the left or right of the midline in 100% of trials. In the low-certainty condition, the target jumped in 60% of the trials and continued to move straight down the midline in 40% of trials (no-jump trials). We further examined the effect of response urgency, by varying the onset of jumps across all trials. Early jumps required a low level of urgency (responses within 450 *ms*), middle jumps required a medium level of urgency (responses within 350 *ms*), and late jumps required a high level of urgency (responses within 250 *ms*).

Using this paradigm, we had two expectations of how jump certainty and response urgency would affect hand responses (Fig. 1A). First, we hypothesized that high-certainty blocks would be associated with earlier (lower reaction time) and faster (higher vigor) hand responses compared to low-certainty blocks. Second, we hypothesized that as the response urgency increased, hand responses would be initiated earlier (lower reaction time) and with greater vigor. However, the goal of the oculomotor system is less clear (Fig. 1B). An anticipatory saccade to the location of interception could provide foveal vision to guide contact between the paddle and the target (Fooken et al., 2021). In this case, we would predict ‘regular’ saccade latencies of ∼200 *ms* (Carpenter & Williams, 1995), and similar modulations of eye reaction times and vigor as observed in hand responses for varying levels of certainty and urgency. Moreover, eye and hand responses would be strongly correlated on a trial-by-trial basis (Fig. 1B). Alternatively, eye responses in this task could be reactive. In this case, we would predict eye responses with similar latencies to express saccades (Fischer & Ramsperger, 1984; Paré & Munoz, 1996). Moreover, we would expect a differential modulation of eye and hand responses by jump certainty and response urgency, and only a weak or no correlation between eye and hand responses (Fig. 1C).

**Figure 1.**
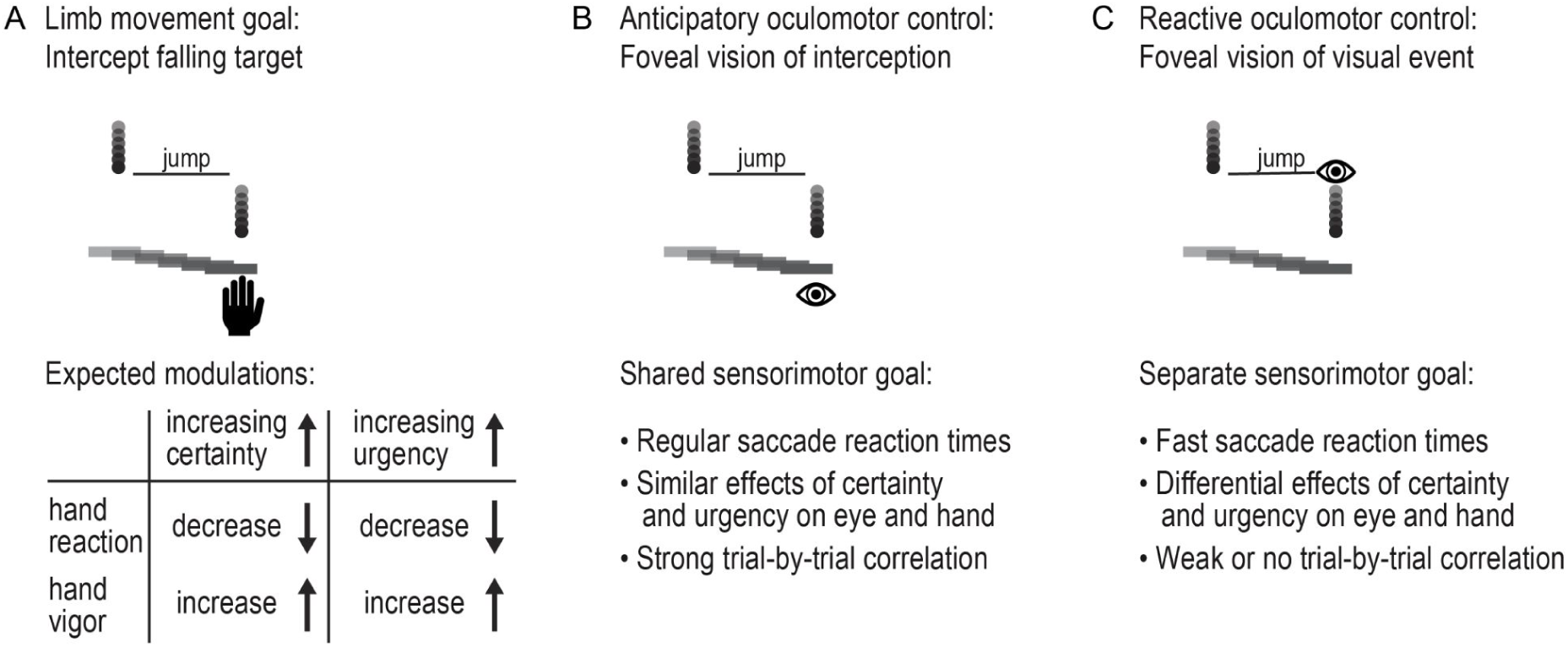
Effects of jump certainty and response urgency on eye and hand responses. (A) Hand movements generated to intercept a falling target are expected to be earlier and faster in high-certainty and more urgent trials. (B) The oculomotor system may share the sensorimotor goal of the upper limb motor system (anticipatory control for interception), in which case the effects of jump certainty and response urgency will be similar. (C) Alternatively, the oculomotor system’s goal may be to detect the visual event (reactive control), in which case eye movements will be differentially affected by changes in jump certainty and response urgency.

## Methods

### Participants

A total of 14 healthy individuals (12 right-handed; 11 females; mean age: 26.5 years; age range: 18-44 years) participated in the experiment. All participants had no self-reported neurological and musculoskeletal impairments and normal or corrected-to-normal vision. The study was approved by the Queen’s University Health Sciences & Affiliated Teaching Hospitals Research Ethics Board (TRAQ #: 6003707) and adheres with the Declaration of Helsinki. Participants gave written informed consent and were compensated with a small honorarium ($10 CAD).

### Apparatus

Experiments were conducted using the Kinarm Endpoint robot (Kinarm^TM^, Kingston, ON, Canada). Participants were seated in a chair supporting their backs, and used one hand to grasp the handle of the robotic manipulandum, which allowed movement along the horizontal plane. Movements made by the participant were tracked by the manipulandum and presented as a cursor on a 32 *inch* vertical display placed 37 *cm* from the participant (Fig. 2A). The mapping between the handle and cursor movement was the same as a standard computer mouse, such that forward and backward movements of the handle moved the cursor up and down, and left and right handle movements moved the cursor left and right. Movements of the handle were recorded at a sampling rate of 1000 *Hz*. The inherent visual display delay was accounted for using the latency reported by the graphics card in the robot computer and the calculated refreshing latency (∼50 *ms*). The position of the participants’ right eye was recorded using a video-based eye tracker (Eyelink, SR Research, ltd., Kanata, ON, Canada) with a sampling rate of 500 *Hz*. A combined chin and forehead rest minimized head movements during the experiment.

**Figure 2.**
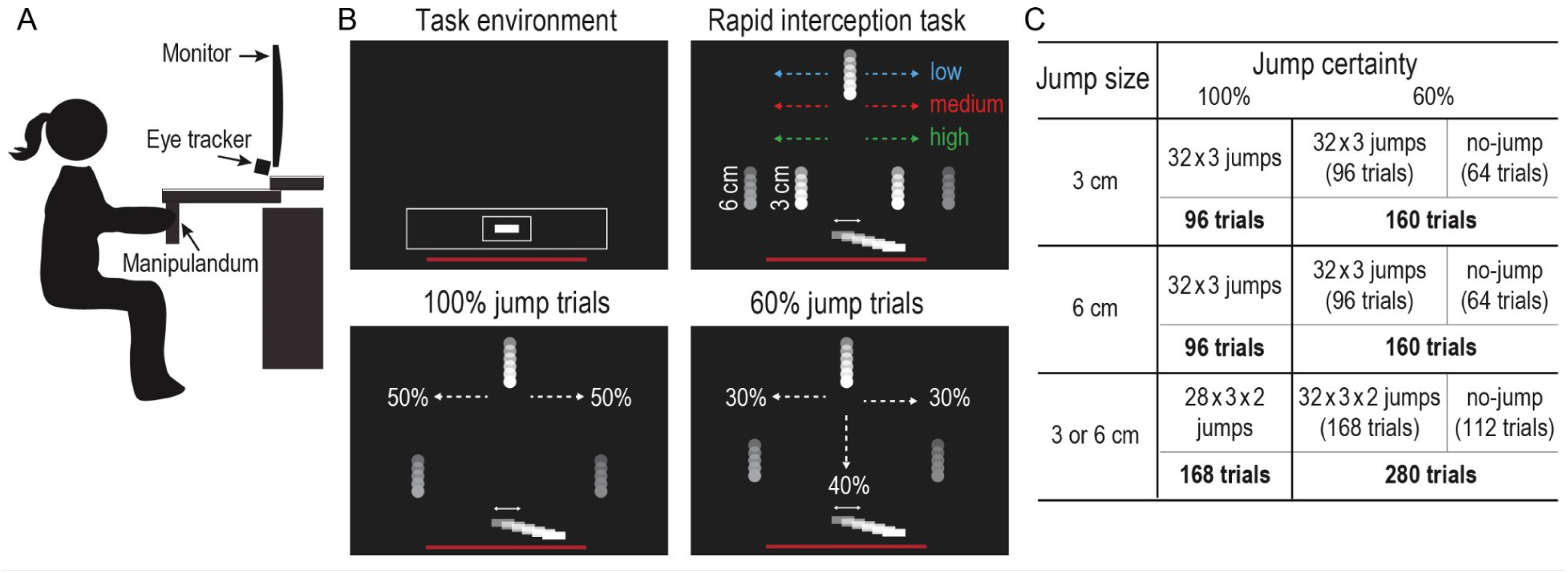
Experimental setup and conditions. (A) Participants moved the handle of a robotic manipulandum in the horizontal plane to intercept moving targets viewed on a vertical screen. (B) Each trial started by showing the task environment (top left). The response urgency was manipulated by changing the position at which the target jumped (top right). The target could jump 3 *cm*, 6 *cm*, or both within a block. In the high-certainty condition (100% jump trials; lower left), the target moved down the midline before it randomly jumped to the left or right side (lower left). In the low-certainty condition, the target could continue to move down the midline (no-jump) in 40% of the trials (lower right). (C) Participants performed 32 or 28 interceptions for each urgency level in different blocks of jump certainty and jump size combinations.

### Experimental Protocol

We modified a previously established interception task to assess the coordination between eye and hand responses (Park et al., 2023). Figure 2B illustrates the rapid interception task, in which participants intercepted a target moving down a vertical screen with a virtual paddle (white rectangle 2 × 0.5 *cm* or 3 × 0.8 *deg*) representing the position of the hand. At the beginning of each trial, participants were shown the task environment, which consisted of the workspace represented by a white box (Workspace, 28 × 3.5 *cm* or 37.1 × 5.4 *deg*) indicating the allowed area of movement. A smaller box (Starting Area, 3.5 × 2 *cm* or 5.4 × 3.1 *deg*) indicated the area that participants had to move their paddle into to begin a trial. A red line representing the bottom limit for intercepting the moving target was displayed to deter individuals from moving below the workspace. After holding the paddle in the starting area for 200 *ms*, the task environment disappeared, and a target (white circle with a radius 0.5 *cm* or 0.8 *deg*) appeared at the top of the screen (22 *cm* or 30.7 *deg* from the centre of the starting area). The target then immediately started to move down the screen at a speed of 25 *cm/s* or 34 *deg/s*. The target either continued to move straight down or could be visually perturbed to the left or right by a same probability. The onset of perturbation was randomized at three different distances from the centre of the starting area (11.25 *cm* or 16.9 *deg*, 8.75 *cm* or 13.3 *deg*, or 6.25 *cm* or 9.6 *deg*). After the perturbation, the target continued to move straight down. Participants had to intercept the target before it reached the bottom of the workspace. They were able to intercept the target with any part of the paddle and in any direction. In case participants left the workspace that was shown at the beginning of the trial (the large white box), the robot applied a step force in the opposite direction of their hand movement in order to push their hand back into the box. On successful interceptions, haptic feedback was applied to the participant’s hand via the robotic manipulandum to indicate success. Subsequent trials began 500 *ms* after the target was intercepted or missed.

To investigate the relationship between eye and hand movements, we manipulated jump certainty and response urgency. We examined certainty by having two groups of blocks of either high-certainty conditions (target jumped in 100% of the trials) or low-certainty (target jumped in 60%). We probed response urgency by varying the onset of visual perturbation. The target was perturbed at three different locations from the top of the workspace, requiring more urgent responses for later target jumps. The highest jump position was 11.25 *cm* (16.9 *deg*) above the Starting Area (low-urgency; ∼450 *ms* to respond), the middle jump position was 8.75 *cm* (13.3 *deg)* above the Starting Area (medium-urgency; ∼350 *ms* to respond), and the lowest jump position was 6.25 *cm* (9.6 *deg*) away (high-urgency; ∼250 *ms* to respond). Response urgency (i.e., jump onset position) was randomized at equal probability across all trials. Further, we examined the magnitude of motor responses by varying the size of the jump (3 *cm* or 4.6 *deg*, 6 *cm* or 9.2 *deg*, or both 3 and 6 *cm* with equal probability). The size of the jump was varied in three separate blocks and for each jump size, we tested both high and low-certainty, resulting in six total blocks (Fig. 2C). Within each certainty condition, the order of blocks with different jump sizes was randomized. Participants completed either three high-certainty blocks, then three low-certainty blocks, or three low-certainty blocks and three high-certainty blocks. There were 960 trials in total, which took participants ∼1 hour to complete, including the breaks.

### Eye and Hand movement analysis

Eye and hand movement data were analyzed offline using custom-made codes in MATLAB R2021a (Mathworks Inc. Natick MA, USA). The x and y positions of the center of the robotic handle were used for hand movement analysis. The position of the handle was filtered using a third order, zero-phase lag, 20 *Hz* Butterworth filter. We analyzed horizontal and vertical interception position, defined as the x and y position of the robotic handle at the time the target first contacted any part of the paddle. We further analyzed hand movement reaction time and vigor. Hand movement reaction time was defined as the difference from the time the visual target jumped to the time of hand movement onset, with hand movement onset being the first moment in time at which the hand velocity was greater than 8% of the peak hand velocity in the current reach. Hand movement vigor was calculated similar to movement vigor previously reported in the literature (Reppert et al., 2015). In each trial, we defined the hand movement amplitude as the absolute furthest distance the hand travelled between the time of target movement onset and the time of target interception, or the time the target left the interception zone in target-miss trials. We then fitted a hyperbolic function for each participant *n* to capture the relationship between movement peak velocity *v*_*n*_ and amplitude *x*

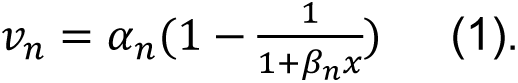

This fit yielded participant-specific parameters of *α*_*n*_ and *β*_*n*_ that were used to calculate the expected movement velocity *v*_*fit*_given each observed movement amplitude. Vigor was defined as a within-participant measure by comparing the actual movement velocity with the expected movement velocity *v*_*n*_/*v*_*fit*_. Thus, a vigor value > 1 indicates that a given movement was faster than expected, and a value < 1 indicates that the movement was slower than the average movement for this amplitude.

The x and y eye positions of the right eye were sampled in screen-centred coordinates, and position signals were filtered using a second-order 15 *Hz* Butterworth low pass filter. We analyzed vertical and horizontal eye positions at selected times throughout the task (time of target movement onset, at the time of target jump, and 250 *ms* after target jump). We further analyzed eye movement accuracy, reaction time, and vigor of the first saccade that occurred at least 50 *ms* following the target jump. Akin to hand movement measures, reaction time was defined as the difference from the time the visual target jumped to the time of saccade onset. Eye movements were labelled as saccades when five consecutive samples of the filtered eye velocity exceeded a fixed velocity criterion of 30 *cm/s*. Saccade on- and offsets were labelled as the time that the sign of the acceleration signal reversed either before eye velocity exceeded (saccade onset) or subceeded (saccade offset) the velocity threshold. To determine eye movement vigor, we followed the same procedure as outlined for hand movement vigor. Saccade accuracy was defined as the x and y distance between eye and target position at the time of saccade offset.

### Statistical Analysis

To evaluate the effects of jump certainty and response urgency on hand and eye movement measures, we calculated the median value for each condition and participant. We then used repeated-measures ANOVAs with an alpha level of 0.05. Post hoc comparisons were done using two-sided, paired *t*-tests with Bonferroni correction. Because we did not have an a priori hypothesis of how jump size (3 or 6 *cm*) would affect selected eye and hand measures, we averaged across jump size unless stated otherwise in the manuscript, Statistical tests were conducted using R (R Core Team, 2022; www.r-project.org).

### Data exclusion

One participant was excluded due to a high number of eye data loss, affecting 47% of their trials. Across the remaining participants’ trials, 2034 out of 13440 (15%) were flagged. Trials were flagged if the eye signal was lost (e.g., due to blinks) in the time window from target movement onset to the time of interception (6.5%) or if the participant would move pre-emptively (8.5%). We labelled hand movements as pre-emptive if the participant’s reaction time was lower than 120 *ms* after the target jumped or if they moved to the opposite side of the jump direction.

## Results

The goal of this experiment was to investigate the coordination of eye and hand responses when intercepting a rapidly moving target that could jump at a variable time to the left or right. In a given block, the target either jumped with high-certainty (100% jump trials) or low-certainty (60% jump trials). In no-jump trials (40% of all trials in the low-certainty condition), the target continued to travel down the midline, and participants were required to keep the paddle at the centre of the screen to intercept the target successfully. We manipulated response urgency by changing the time at which the target jumped (see Fig. 2B).

### Patterns of eye and hand movements are similar in jump and no-jump trials

The rapid interception task elicited a combination of tracking and saccadic eye movements. The upper panels in Figure 3 show the eye, hand, and target position of a representative high-certainty trial in a screen-centred reference frame (Fig. 3A) and across time (Fig. 3B). Eye, hand, and target velocity of the same trial are also shown across time (Fig. 3C). In this trial, the participant looked at the centre of the screen and ‘waited’ for the target to move down the midline. The participant briefly tracked the target with smooth pursuit eye movements before initiating a reactive saccade towards the target after it jumped to the right.

**Figure 3.**
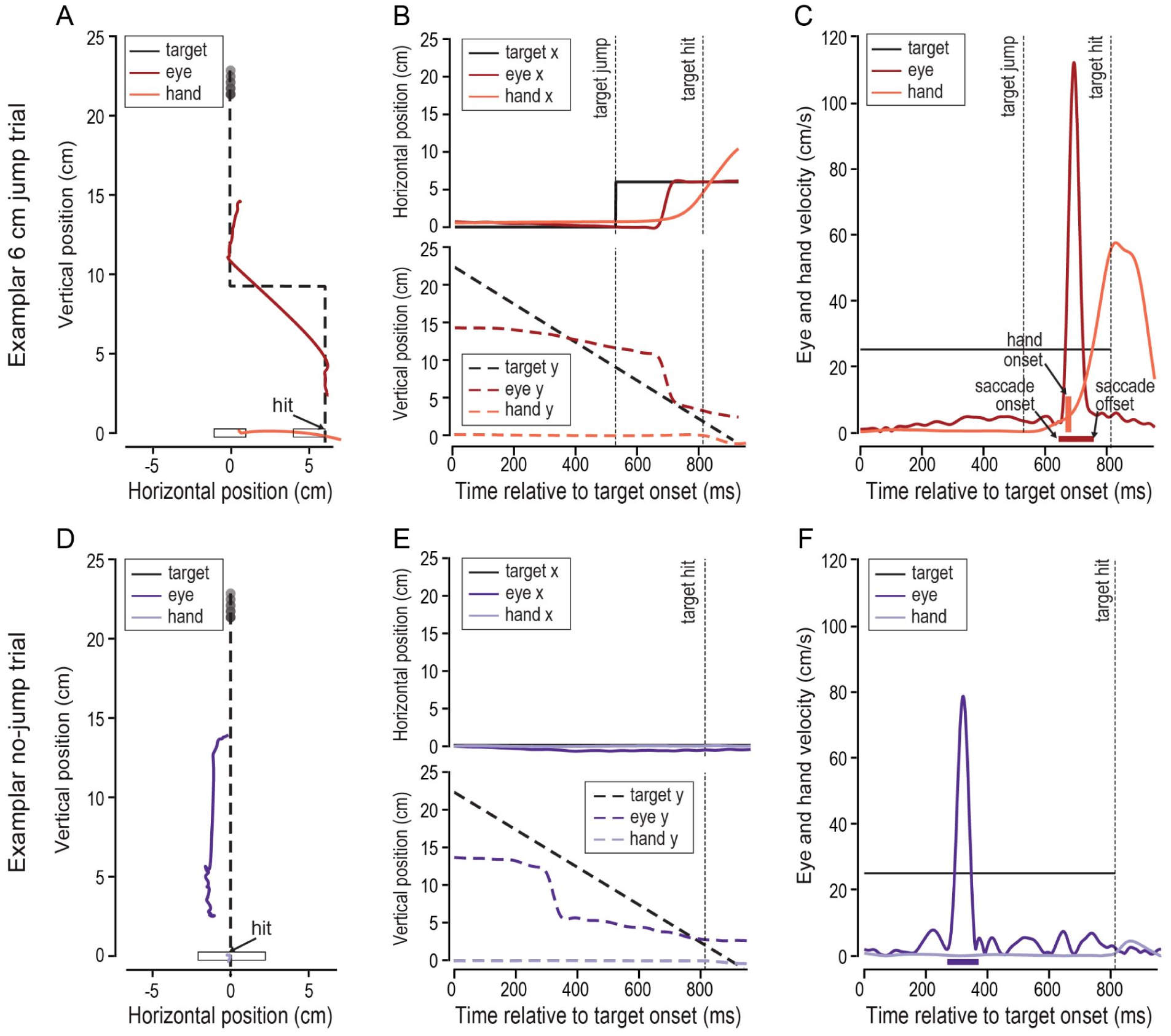
Eye and hand movement responses in the rapid interception task. (A) Eye (dark red), hand (light red), and target (black) position in screen-centred coordinates in a representative jump trial. (B) Horizontal eye, hand, and target position (top panel) and vertical eye, hand, and target position (bottom panel) across time. The time of target jump and target interception are indicated by dashed vertical lines. (C) Eye, hand, and target velocity across time. Saccade onset and offset of the targeting saccade following the target jump are indicated by arrows and the thick horizontal bar. Hand movement onset is indicated by the thick vertical bar. (D) Eye (dark purple), hand (light purple), and target (black) position in screen-centred coordinates in a representative no-jump trial. (E) Horizontal eye, hand, and target position (top panel) and vertical eye, hand, and target position (bottom panel) across time. The time of the target interception is indicated by the dashed vertical line. (F) Eye, hand, and target velocity across time.

The lower panels in Figure 3 show the eye, hand, and target position of a representative low-certainty trial—in which the target did not jump—in a screen-centred reference frame (Fig. 3D) and across time (Fig. 3E). Eye, hand, and the target velocity of the same trial are also shown across time (Fig. 3F). Similar to the perturbation trial, the participant looked at the centre of the screen at the beginning of the trial. The participant tracked the target with smooth pursuit and a catch-up saccade until it was intercepted by keeping the paddle on the midline.

Across all trials and participants, and in both certainty conditions (including jump and no-jump trials), we found a consistent pattern of eye movements. In the time window from target movement onset to the time of target jump or target interception for no-jump trials, participants made no saccades in 42% of the trials, made a single saccade in 47% of the trials, and elicited more than one saccade in 11% of the trials. Eye velocity in the same time window was on average 7.58±0.97 *cm/s* (mean eye velocity and standard error across participants), which was much lower than the target velocity of 25 *cm/s,* indicating that participants did not smoothly pursue the moving target prior to target jump. Following a target jump (in the jump trials), participants initiated a reactive saccade in 93% of all trials. Of note, in a majority of these trials (96%) participants did not make another saccade before they intercepted the moving target. The described eye movement patterns show that whereas eye movements prior to target jump were quite variable (short periods of pursuit, catch-up saccades, or fixation), a single reactive saccade was elicited following target jump in almost every trial.

### Eye movement position is modulated by the response urgency

Figure 4 illustrates the overall gaze pattern observed in our experiment. Figure 4A shows the gaze position (from target movement onset to target interception) of all participants (thin lines) in screen-centred coordinates averaged across high-certainty 3 *cm* (top) and 6 *cm* (bottom) jump trials, respectively. The gaze position averaged across participants at different urgency levels is indicated by thick blue (low-urgency), red (medium-urgency), and green (high-urgency) lines. Figure 4E shows the corresponding screen-centred gaze position in low-certainty trials. To further describe the observed gaze pattern, we compared participants’ eye position at three distinct time points: (1) at the time of target movement onset (Fig. 4B, F), (2) at the time of target jump (Fig. 4C, G), and (3) 250 *ms* after the time of jump (Fig. 4D, H). We chose 250 *ms* to allow sufficient time for saccades to land following the target jump.

**Figure 4.**
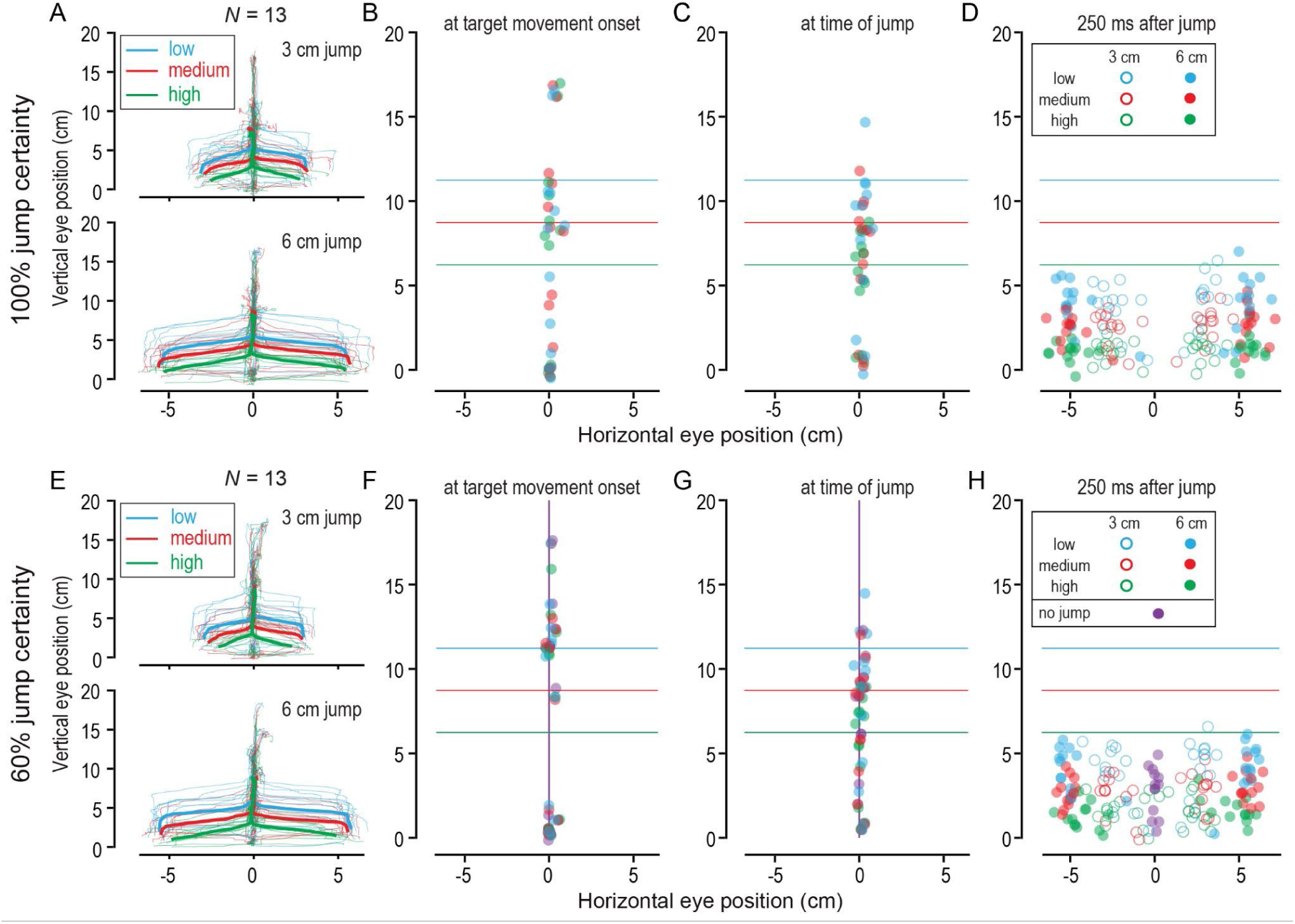
Eye responses in rapid interception task. (A) Gaze position in 100% certainty trials averaged across low (blue), medium (red), and high (green) urgency trials of each participant (thin lines) and averaged across the group (thick lines) separately plotted for the 3 *cm* (top) and 6 *cm* (bottom) target jump size conditions. (B-D) The average 2D eye position is shown for each participant and each urgency level at the time of the target movement onset (B), the time of the target jump (C), and 250 *ms* after the target jump (D). (E-H) Corresponding plots for the 60% jump certainty condition. Eye positions in no-jump trials are indicated in purple.

We first compared the effect of certainty (high vs. low) and urgency (low, medium, high) on vertical eye position at the three selected time points using a repeated-measures 2×3 ANOVA. We found no effect of certainty, urgency, and no interaction on vertical eye position at the time of target movement onset (*F* < 1.7; *p* > 0.2; *η* < 0.13). We also found no effect of certainty and no interaction on vertical eye position at the time of the target jump (*F* < 0.7; *p* > 0.5; *η* < 0.06). We found an effect of urgency on vertical eye position at the time of target jump (*F_2,24_*= 20.72; *p* < 0.001; *η* = 0.63). Finally, we did not find an effect of jump certainty (*F* < 1.5; *p* > 0.2; *η* < 0.12) on vertical eye position 250 *ms* after target jump, but a significant effect of urgency (*F_2,24_* = 91.82; *p* < 0.001; *η* = 0.88) and a significant interaction between jump certainty and response urgency (*F_2,24_*= 6.52; *p* = 0.005; *η* = 0.35). Taken together, these results indicate that vertical eye position was significantly affected by target jump times with vertical eye position being lower as the urgency level increased. Vertical eye position was not affected by jump certainty. As illustrated by each participant’s gaze position (Figs. 4A, E), the between-participants variability was high at target movement onset (∼6 *cm*), but participants converged to similar vertical gaze positions following the target jump.

Horizontal gaze positions were near the midline until after the target jumped (Figs. 4 B-C, F-G), and horizontal gaze remained at the midline during trials where the target did not jump (purple dots in Figs. 4 F-H). Following the target jump, the horizontal eye position scaled with the size of the target jump, landing on average 2.65±0.16 *cm* away from the midline in 3 *cm* jump trials and 5.29±0.09 *cm* away from the midline in 6 *cm* jump trials.

Finally, we investigated the accuracy of the first saccade that occurred after target jump. Whereas horizontal saccade accuracy was unaffected by jump certainty and response urgency, vertical saccade accuracy was affected by jump certainty (*F_1,12_* = 4.85; *p* = 0.048; *η* = 0.29) and response urgency (*F_2,24_* = 123.92; *p* < 0.001; *η* = 0.91). Post hoc comparison showed that participants were on average very accurate in high-certainty blocks (vertical saccade error: −0.08±0.32 *cm*), but tended to land above, or behind, the actual target position in low-certainty blocks (vertical saccade error: 0.27±0.32 *cm*). Moreover, saccades were most accurate in medium-urgency trials (vertical saccade error: 0.05±0.19 *cm*). In low-urgency trials, saccades tended to land below, or ahead, of the moving target position (vertical saccade error: −0.94±0.26 *cm*), and in high-urgency trials saccades tended to land above, or behind, the moving target position (vertical saccade error: 1.19±0.16 *cm*).

### Interception accuracy decreases with uncertainty and increasing urgency

Figure 5 illustrates the overall hand movements observed in our experiment. Figures 5A and B show the horizontal hand position (upper panels) and absolute hand velocity (lower panels) during high-certainty 6 *cm* jump trials for an exemplar participant and the average of all participants, respectively. The corresponding plots of the horizontal hand position and velocity during 6 *cm* low-certainty trials are shown in Figures 5 D and E. In these plots, hand movements are averaged across different levels of urgency as indicated by colour. Because the moving target could be intercepted with any part of the paddle, there is a 4 *cm* wide region and a limited time window to intercept the falling target successfully (see grey-shaded regions and horizontal, coloured lines in panels of Fig. 5A-B, D-E). We found that the onset of participants’ hand responses scaled with response urgency.

**Figure 5.**
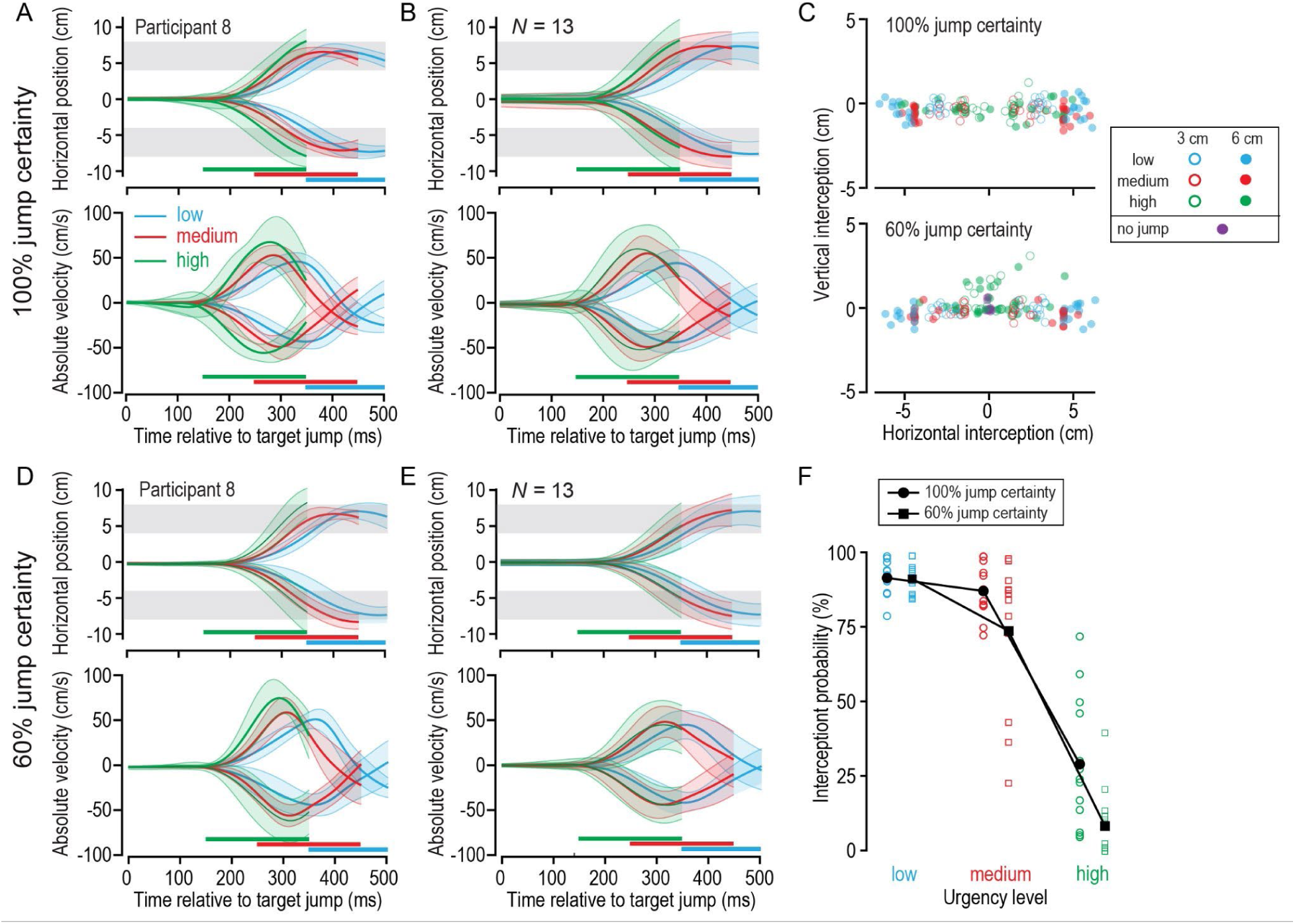
Hand responses in the rapid interception task. (A) Horizontal hand position (top) and velocity (bottom) across time for low (blue), medium (red), and high (green) urgency trials shown for an exemplar participant in the high-certainty condition. The grey-shaded areas represent the region in which the paddle could intercept the target. The horizontal coloured bars indicate the time period in which the target had to be intercepted for each urgency level. (B) Corresponding plots averaged across all participants in the high-certainty condition. (C) Hand position at the time of interception or when the missed target left the screen in the high-certainty (top) and low-certainty (bottom) conditions. Open circles indicate 3 *cm* jump trials and filled circles indicate 6 *cm* jump trials. Colour denotes urgency level, and purple indicates trials in which the target did not jump. (D-E) Plots of horizontal hand position and velocity across time for an exemplary participant (D) and averaged across all participants (E) in the low-certainty condition. (F) Probability of intercepting the moving target following a target jump in the high-certainty (circles) and low-certainty (squares) jump conditions and across urgency levels.

To compare performance across conditions, we analyzed hand position at the time of the target interception and the probability of successfully intercepting the falling target across conditions. Figure 5C shows for each certainty condition, urgency level, and jump size, the average screen-centred hand position at the time of the target interception or when the missed target left the screen. Overall, hand movement scaled with the size of the jump, and participants intercepted the target on average 2.02±0.10 *cm* away from the midline in 3 *cm* jump trials and 3.95±0.15 *cm* away from the midline in 6 *cm* jump trials. The fact that participants undershot the actual target position indicates that they tended to contact the falling target with the outer side of the paddle.

We used a repeated-measures 2×3 ANOVA to quantify the effect of certainty and urgency on horizontal interception position. We found an effect of jump certainty (*F_1,12_* = 30.55; *p* < 0.001; *η* = 0.72), response urgency (*F_2,24_* = 193.09; *p* < 0.001; *η* = 0.94), and a significant interaction (*F_2,24_* = 4.18; *p* = 0.028; *η* = 0.26). The strong effect of response urgency on horizontal interception position reflects the shorter time available for intercepting the perturbed target (see Figs. 5 A-B, D-E). Participants intercepted the target with the centre of the paddle in low-urgency trials, but intercepted the target with the outer edge of the paddle in medium and high-urgency trials. On average, participants tended to move the paddle slightly downwards to intercept the falling object (vertical interception position: −0.27±0.06 *cm*). Interestingly, we observed that in the low-certainty and high-urgency conditions, participants tended to move the paddle upwards (vertical interception position: 0.34±0.22 *cm*), as if they expected the target to stay in the middle (see green dots above the centre in the lower panel of Fig. 5C).

Figure 5F shows the averaged probability of intercepting the falling target across task conditions. Individual colour-coded symbols represent the median response value of each participant, and larger black-coloured symbols (connected by black lines) represent the group means. We found an effect of jump certainty (*F_1,12_*= 24.27; *p* < 0.001; *η* = 0.67), response urgency (*F_2,24_* = 194.12; *p* < 0.001; *η* = 0.94), and a significant interaction effect (*F_2,24_* = 5.73; *p* = 0.009; *η* = 0.32). Whereas participants successfully intercepted a majority of targets in low-urgency trials (high-certainty: 91.7±1.5%; low-certainty: 91.4±1.4%) and medium-urgency trials (high-certainty: 87.3±2.4%; low-certainty: 73.9±6.7%), they only intercepted about a third of targets in high-certainty, high-urgency trials (29.2±6.0%) and only a few targets in low-certainty, high-urgency trials (8.5±3.2%).

### Differential effects of certainty and urgency on eye and hand responses

Figure 6 shows the eye and hand reaction times, vigor, and trial-by-trial correlation between eye and hand responses for the two certainty conditions and three urgency levels. To directly compare eye and hand responses, only trials in which a saccade was made (93% of all trials) were included in this analysis. To test the effect of jump certainty and response urgency on eye and hand responses, we used four separate repeated-measures 2×3 ANOVA. Hand responses were on average initiated 188.5±3.4 *ms* after the target jump. We found an effect of jump certainty (*F_1,12_* = 27.04; *p* < 0.001; *η* = 0.69) and response urgency (*F_2,24_* = 13.86; *p* < 0.001; *η* = 0.54) on hand reaction time, and no significant interaction (Fig. 6A). A post hoc comparison of the two certainty conditions confirmed that participants on average initiated their hand responses 14 *ms* earlier in the high-certainty blocks compared to the low-certainty blocks (*t*(38) = 6.8; *p_adjust_* < 0.001; *d* = 1.1). Compared to low-urgency trials, hand responses were initiated 18 *ms* earlier in medium-urgency trials (*t*(25) = 7.4; *p_adjust_* < 0.001; *d* = 1.4) and 19 *ms* earlier in high-urgency trials (*t*(25) = 4.3; *p_adjust_*< 0.001; *d* = 0.8).

**Figure 6.**
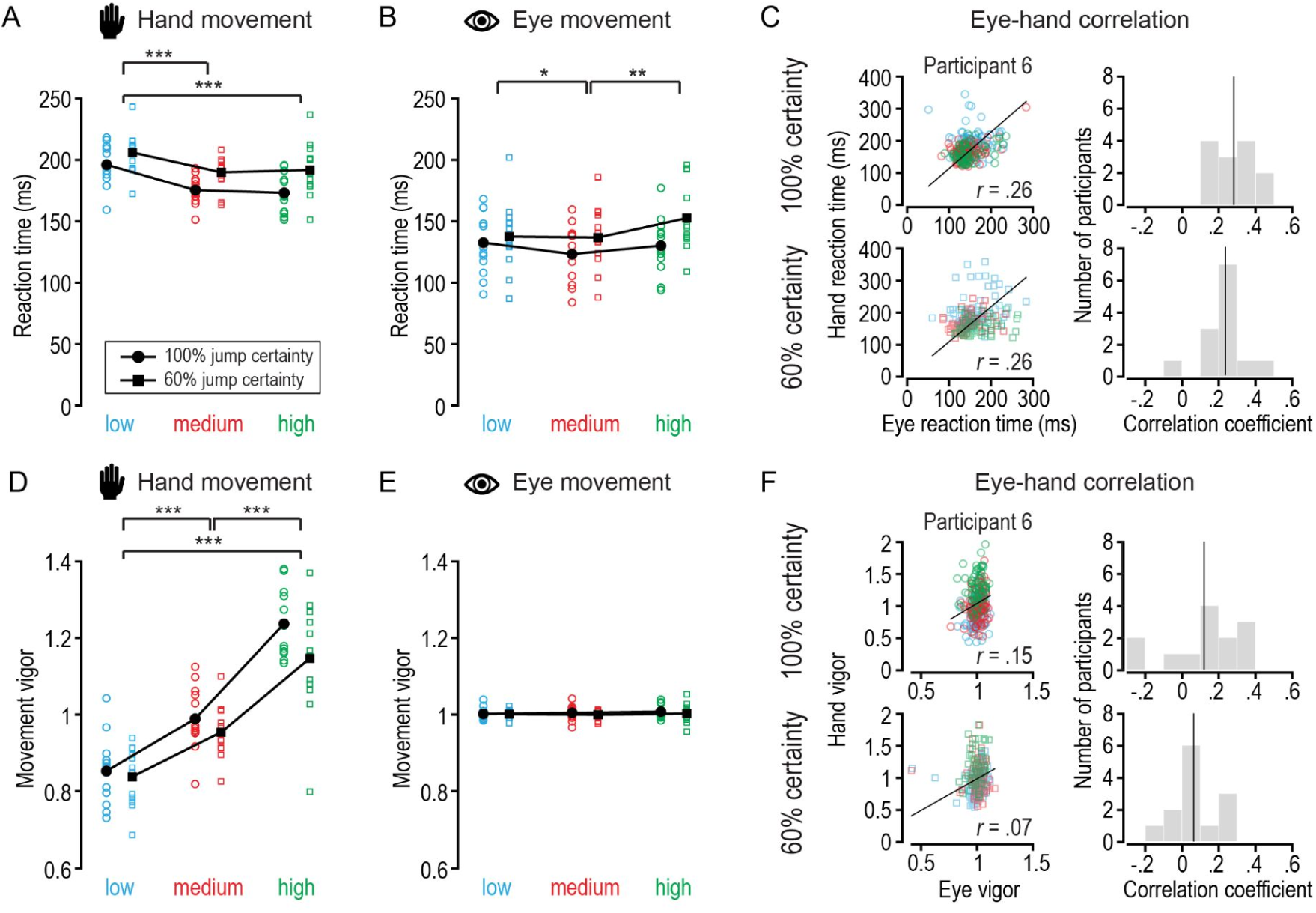
Eye and hand responses across task conditions. (A) Hand movement reaction times for different task conditions. Single symbols indicate median reaction times for each participant for high-certainty jumps (circles) and low-certainty jumps (squares), and separate for low (blue), medium (red), and high (green) urgency trials. (B) Corresponding eye reaction times. (C) Trial-by-trial correlation between eye and hand responses for high-certainty (top left panel) and low-certainty (bottom left panel) trials for a representative single participant, and histograms of the correlation coefficients of all participants in the high-certainty (top right panel) and low-certainty (bottom right panel) condition. (D-F) Corresponding results for hand and eye vigor.

Eye movements preceded hand movements by ∼50 *ms* and were on average initiated 135.4±6.3 *ms* after the target jump. We found an effect of jump certainty (*F_1,12_* = 11.72; *p* = 0.005; *η* = 0.49) and response urgency (*F_2,24_*= 5.27; *p* = 0.01; *η* = 0.31) on eye reaction time, and a significant interaction (*F_2,24_* = 13.48; *p* < 0.001; *η* = 0.53; Fig. 6B). Posthoc comparison of the two certainty conditions confirmed that participants on average initiated their eye movements 14 *ms* earlier in the high-certainty blocks compared to low-certainty blocks (*t*(38) = 5.0; *p_adjust_* < 0.001; *d* = 0.8). Compared to medium-urgency trials, eye responses were initiated 5 *ms* later in low-urgency trials (*t*(25) = 2.8; *p_adjust_* = 0.03; *d* = 0.6) and 11 *ms* later in high-urgency trials (*t*(25) = 4.1; *p* = 0.001; *d* = 0.8). Thus, whereas hand reaction times were systematically faster for high-urgency levels, eye reaction times were differentially affected. On a trial-by-trial level, eye and hand reaction times were on average weakly correlated with a mean correlation of *r =* 0.28 in high-certainty trials and *r =* 0.24 in low-certainty trials (Fig. 6C).

We found an effect of response urgency (*F_2,24_* = 68.71; *p* < 0.001; *η* = 0.85) on hand vigor and a significant interaction between jump certainty and response urgency (*F_2,24_* = 3.42; *p* = 0.049; *η* = 0.22). However, jump certainty did not systematically affect hand vigor (*F_1,12_* < 2.2; *p* > 0.1; *η* < 0.2; Fig. 6D). Post hoc comparison of different urgency levels confirmed that participants responded more vigorously with increasing urgency. Specifically, hand vigor was greater in medium (*t*(25) = 8.7; *p_adjust_* < 0.001; *d* = 1.7) and high-urgency trials compared to low-urgency trials (*t*(25) = 11.3; *p_adjust_* < 0.001; *d* = 2.2), and hand vigor was also greater in high compared to medium-urgency trials (*t*(25) = 9.4; *p_adjust_*< 0.001; *d* = 1.9).

Finally, we found no effect of jump certainty or response urgency on eye vigor (all *F* < 5.2; *p >* 0.5; *η* < 0.05; Fig. 6E). On a trial-by-trial level, eye and hand vigor were on average very weakly correlated, with a mean correlation of *r =* 0.12 in high-certainty jump trials and *r =* 0.06 in low-certainty jump trials (Fig. 6F). Thus, whereas hand movements became more vigorous as the urgency level increased, eye vigor was unaffected and not related to the hand response.

### Jump size differentially modulates eye and hand movement reaction times

We additionally investigated the effect of block type (constant jump size or mixed jump size) and jump size (3 or 6 *cm*) on eye and hand reaction times and vigor. Hand reaction time was not affected by block type, but we found an effect of jump size (*F_1,12_* = 5.68; *p* = 0.03; *η* = 0.32) and a significant interaction (*F_1,12_* = 13.31; *p* = 0.003; *η* = 0.53). Post hoc comparison showed that participants responded on average 7.3 *ms* earlier to 3 *cm* jumps compared to 6 *cm* jumps in the mixed blocks (*t*(12) = 7.7; *p_adjust_* < 0.001; *d* = 2.1), but no difference in hand reaction time was found between 3 *cm* and 6 *cm* jump blocks. Eye reaction time was affected by block type (*F_1,12_*= 12.77; *p* = 0.004; *η* = 0.52) and jump size (*F_1,12_*= 43.40; *p* < 0.001; *η* = 0.78) with no significant interaction. Specifically, participants initiated eye movements on average 5 ms earlier in blocks with constant jump size compared to mixed jump size, and 13.5 *ms* earlier in 6 *cm* compared to 3 *cm* jump trials. These results indicate that eye and hand reaction times were differentially affected, with the hands responding earlier to small target jumps and the eyes responding earlier to large target jumps.

Hand vigor was not affected by block type, but we found an effect of jump size (*F_1,12_*= 8.35; *p* < 0.001; *η* = 0.87) and a significant interaction (*F_1,12_* = 23.08; *p* < 0.001; *η* = 0.87). Post hoc comparison showed that participants responded more vigorously in 6 *cm* compared to 3 *cm* jump trials in the 3 *cm* and 6 *cm* blocks (*t* = 8.1; *p_adjust_* < 0.001; *d =* 2.2) and mixed jump size blocks (*t* = 8.9; *p_adjust_* < 0.001; *d =* 2.5). Eye vigor was not affected by block type but by jump size (*F_1,12_* = 13..36; *p* < 0.003; *η* = 0.53), with participants responding more vigorously in 6 *cm* compared to 3 *cm* jump trials. There was no significant interaction. These results indicate that the strength of the response (i.e., the vigor) was affected similarly for the eye and hand movement system.

## Discussion

In this study, we investigated the coordination of eye and hand responses during a rapid interception task. We found that eye and hand responses were differentially affected by various levels of jump certainty and response urgency (Fig. 7A). Low-certainty conditions caused a delay in both eye and hand reaction times compared to high-certainty conditions. However, high-urgency conditions systematically led to earlier and more vigorous hand responses, while eye responses were largely unaffected by changes in response urgency.

**Figure 7.**
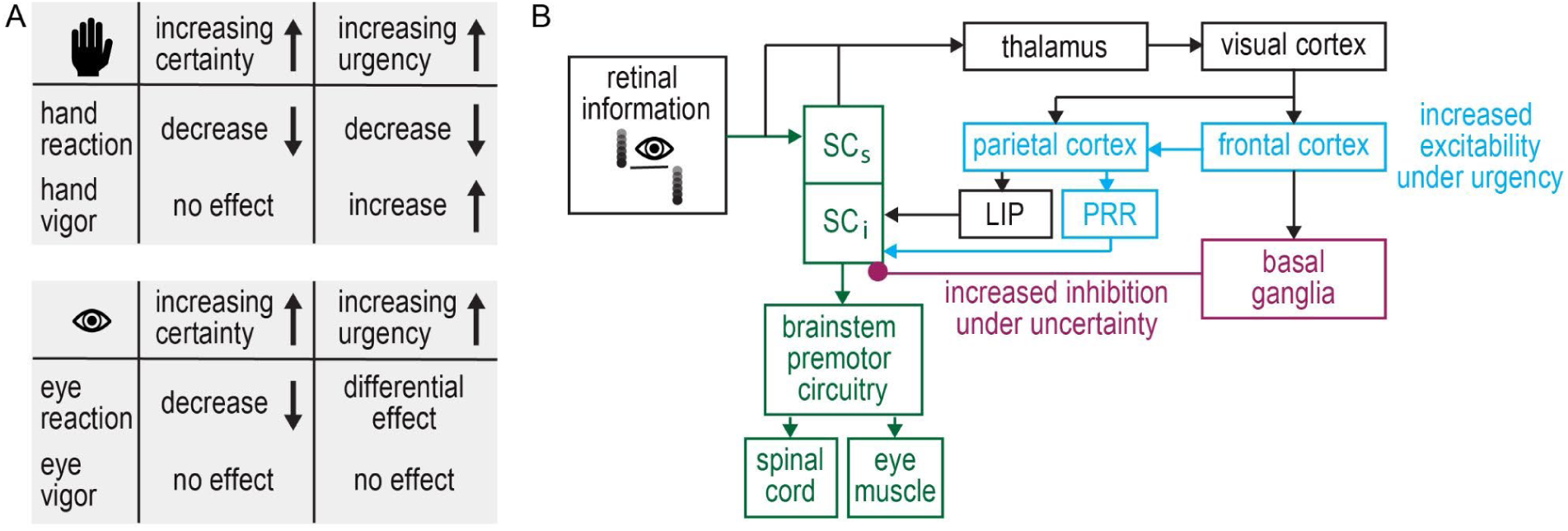
Summary of results and proposed neural modulations. (A) Effects of jump certainty and response urgency on hand (top) and eye (bottom) reaction time and vigor. (B) Simplified visuomotor circuit, highlighting areas involved in rapid responses. Retinal information is directly projected to the superficial layers of the superior colliculus (SC_s_) and transferred to the intermediate layers (SC_i_) that are involved in limb- and oculomotor control (Green). Retinal information also travels through the thalamus to the visual cortex and higher cortical areas. The parietal reach region (PRR) and the lateral intraparietal area (LIP) may drive signals to generate limb and eye responses in parallel.

### Eye-hand coordination depends on task demands

Goal-directed actions in natural environments require an integration of bottom-up sensory information and top-down cognitive goals (Fooken et al., 2023; Scott, 2016). Past research has shown that when reaching towards and manipulating stationary objects, eye and hand movements are highly coordinated in space and time (de Brouwer et al., 2021). When reaching to stationary visual targets, eye and hand reaction times are generally moderately correlated (Fisk & Goodale, 1985; Prablanc et al., 1979)). Interestingly, the correlation between eye and hand reaction time is weaker when the visual stimulus elicits rapid eye responses, such as in the gap paradigm, in which the initial fixation target is extinguished before the saccade target appears (Saslow, 1967). Moreover, the correlation between eye and hand responses is stronger when the task requires a cognitive component, such as memory-guided movements or movements to the opposite side of the cued target (Gribble et al., 2002; Sailer et al., 2000). The rapid interception task used in this study elicited rapid eye and hand responses to visual perturbations. Similar to stationary reaching, we found that rapid eye and hand responses to intercept a perturbed moving target were only weakly correlated (*r* ≈ 0.25). Taken together, these results indicate that correlation between eye and hand movements depends on the visual information available and the cognitive task demands (Frens & Erkelens, 1991).

When intercepting or manually tracking moving objects, observers naturally keep their eyes on the target until they hit or catch it (Cesqui et al., 2015; Coudiere & Danion, 2024; Mrotek & Soechting, 2007). To compensate for high target speeds, the eyes typically track moving objects with a combination of smooth pursuit and saccadic eye movements (Binaee & Diaz, 2019; Fooken et al., 2016; Goettker, Brenner, et al., 2019). In interception tasks that involve distinct visual predictions, such as a target bounce or spatially restricted interception zones, observers make anticipatory saccades to the future target location (Diaz et al., 2013; Fooken & Spering, 2020; Land & McLeod, 2000; Mann et al., 2019). Whereas better object tracking is correlated with higher interception accuracy when target motion is unpredictable and uncertain, eye-hand coordination is more flexible in situations of high motion predictability and visual certainty (Fooken et al., 2021). In our study, we found that participants did not reliably track the moving target prior to the target jump (Fig. 4). Eye position between the time of target movement onset and the time of target jump varied extensively between participants, and the average eye velocity in the same time window was well below target velocity. These results indicate that participants did not attempt to keep their eyes on the moving target, but instead kept their eyes relatively still to detect the target jump.

These results support the hypothesis that the oculomotor response was driven by the salient sensory event rather than by planning an anticipatory saccade to the future interception location (Fig. 1B-C). Participants initiated saccades towards the location of the jumped target with latencies of ∼135 *ms*, which is slightly slower than human express saccades elicited in the gap paradigm (Fischer & Ramsperger, 1984) but faster than ‘regular’ visually-driven saccades (Irving et al., 2006). Previous work has shown that saccade reaction times to moving targets depend on the visual features of the target, such as target velocity (Gellman & Carl, 1991; Ron et al., 1989) or target contrast (Goettker, Braun, et al., 2019). Importantly, these saccades are initiated with latencies of at least 170 *ms* and are able to compensate for target motion during saccade planning and execution (Engel et al., 1999; Etchells et al., 2010; Gellman & Carl, 1991; Goettker et al., 2019; Ron et al., 1989). Here, we found that saccades in response to target jumps landed ahead of the moving target in low-urgency trials and behind the moving target in high-urgency trials. These results indicate that saccades were reactive, with limited time for anticipating the interception location (Findlay, 1983; Robinson, 2022). Interestingly, we found that participants did not make a second saccade to the interception location, even when participants had up to 500 *ms* to intercept following the target jump (low-urgency condition). This finding is in line with previous research showing that participants tend to suppress saccades shortly before intercepting moving objects (Goettker et al., 2019; Mrotek & Soechting, 2007). Overall, our results demonstrate that eye and hand responses are generally coordinated—but not necessarily correlated—to accomplish rapid goal-directed interceptions.

### Rapid visuomotor responses depend on target predictability

When reaching to or looking at visual targets, humans are able to quickly react to sudden changes in the target position, a phenomenon that has been termed express visuomotor or stimulus-locked responses (Goonetilleke et al., 2015; Gu et al., 2016; Pruszynski et al., 2010; Wood et al., 2015). These fast orienting responses can be behaviourally measured across different movement systems, including the oculomotor and limb motor system. Previous work has shown that humans reliably elicit express visuomotor responses when intercepting a vertically moving target that is displaced behind an occluder (Kozak et al., 2020), and that these responses depend on temporal predictability (Contemori et al., 2021; Jakobs et al., 2009). Here, we use a similar experimental paradigm to investigate coordinated eye and hand responses to moving targets. We found that both eye and hand responses following the target jump were initiated earlier in trials in which a target perturbation was highly predictable (high-certainty blocks). Although the possible timings of the perturbation did not change across the experiment, participants had to inhibit a movement in 40% of the low-certainty blocks. These results suggest that participants’ state of response readiness was reduced in the low-certainty condition, similar to an observed decrease in eye reaction time when catch trials are included in the gap paradigm (Kingstone & Klein, 1993; Paré & Munoz, 1996). Response readiness is also increased when targets rapidly move to a new location rather than being instantaneously displaced from one location to another (Reschechtko et al., 2023), indicating that continuous motion prediction is important for rapid visuomotor responses.

In dynamic environments, humans are able to rapidly integrate contextual information to generate motor corrections (Kalidindi & Crévécoeur, 2023). The onset and strength of express visuomotor responses following visual perturbations depend not only on visual target features, such as stimulus luminance, orientation, or spatial frequency (Kozak et al., 2019; Marino et al., 2012; Veerman et al., 2008), but also on the behavioural context, such as target shape, texture, colour, or contextual cues (Contemori et al., 2022; Cross et al., 2019; Dimitriou et al., 2013; Veerman et al., 2008). Moreover, following mechanical perturbations, muscle responses and movement kinematics scale with the time available to respond (Crévécoeur et al., 2013; Poscente et al., 2021). Similarly, we found that hand responses were initiated earlier and more vigorously with increasing urgency, in their respective certainty conditions. The fact that we did not manipulate visual features at the time of perturbation suggests that hand response modulations across urgency levels were driven by the behavioural rather than visual context.

In contrast to the systematic effect of urgency on hand responses, we found that eye reaction time was lowest for medium compared to low and high-urgency trials, and that eye vigor was generally unaffected. Previous research has shown that saccade vigor is indicative of the subjective and economic value of visual targets in value-based decision-making tasks (Korbisch et al., 2022; Reppert et al., 2015; Shadmehr et al., 2019; Yoon et al., 2020). Moreover, saccade vigor is increased when observers have to make a perceptual decision about a moving target in a manual interception task (Barany et al., 2020). Perceptual decisions in urgent visuomotor tasks requiring saccade choices between two possible target locations have been shown to occur after an initial sensorimotor processing time of 90-180 *ms* (Salinas et al., 2019; Seideman et al., 2018; Stanford et al., 2010; Stanford & Salinas, 2021). Taken together, our results suggest that eye vigor is indicative of perceptual decisions and might be computed after an initial bottom-up sensory processing period.

### Neural mechanisms underlying rapid visuomotor responses

To successfully respond to a visual or mechanical perturbation within a few hundred milliseconds, the brain must rapidly transform visual input into motor output, a process that has been shown to involve subcortical circuits. In both the oculomotor and limb motor systems, the superior colliculus (SC)—a midbrain structure that interfaces sensory and premotor circuits—is involved in generating rapid orienting responses (Boehnke & Munoz, 2008; Cooper & McPeek, 2021; Corneil & Munoz, 2014; Gandhi & Katnani, 2011). In the oculomotor system, it has been shown that reduced activity of fixation-related neurons in the SC is linked to the generation of express saccades in the gap paradigm (Dorris et al., 1997, 2002; Marino et al., 2015; Sparks et al., 2000). In the limb motor system, muscle responses to visual or mechanical perturbations can be detected as early as 90 *ms* (Corneil et al., 2004; Gu et al., 2016; Pruszynski et al., 2010; Wood et al., 2015), and rapid muscle activity is highly correlated with neural activity in collicular reach cells (Philipp & Hoffmann, 2014; Stuphorn et al., 1999; Werner et al., 1997). Moreover, when intercepting a moving target that abruptly shifts its position, muscle activity of human participants is modulated 90-110 *ms* after the perturbation, indicating that the speed of these rapid responses would likely involve subcortical pathways (Perfiliev et al., 2010). We believe that the rapid visuomotor responses observed in our study involve pathways through the superior colliculus-brainstem loop.

Extensive work on visually guided eye movements has highlighted the subcortical mechanisms contributing to saccade generation and suppression (for a review, see Coe & Munoz, 2017). We propose that in our task, two factors may modulate eye and hand responses: 1) a reduced inhibition and 2) an increased excitability of the intermediate layers of the SC. Our observation that both eye and hand responses occurred earlier in high-certainty compared to low-certainty blocks suggests a common mechanism modulating visuomotor responses. We speculate that a change in certainty about an upcoming target jump affected inhibitory control from the basal ganglia to the SC (Fig. 7B). Whereas inhibitory mechanisms were upregulated in low-certainty blocks and visuomotor responses had to be withheld in 40% of the trials, inhibition was reduced in high-certainty blocks. This potential regulation of inhibition affected both the oculomotor and limb motor systems similarly.

Our observation that eye and hand responses were differentially affected by urgency suggests that shared sensory information is used differently by the oculomotor and limb motor systems. We speculate that increasing urgency led to an increase in excitability along the limb motor pathway, possibly driven by top-down cortical processes (Fig. 7B). Previous work on the neural control of eye-hand coordination has elucidated that both movement systems are controlled by shared early sensory and visuomotor processing areas (Battaglia-Mayer et al., 2003; Crawford et al., 2004; Dean et al., 2012; Hwang et al., 2014; Vesia & Crawford, 2012). More recently, the idea that the oculomotor and limb systems use the same task-specific sensory information, but operate in parallel to attain task goals has been brought forward (Kang et al., 2024). Our results support the idea of a parallel functional organisation of eye-hand coordination stem that relies on shared sensory information but may serve separate behavioural goals (Fig. 1).

### Eye-hand coordination depends on spatiotemporal task constraints

When controlling goal-directed actions, the human motor system must compensate for inherent delays that arise from transmitting and processing sensory input through neural pathways. For example, to accurately plan a saccade to a moving target, the oculomotor system has to rely on internal information about the trajectory and duration of the eye movement to compensate for the object motion (Schlag & Schlag-Rey, 2002). When planning a hand movement to a moving target, there is a similar delay of ∼100 *ms* to transform visual information into the limb motor command (Brenner & Smeets, 1997). However, unlike saccades, hand movements can be corrected online after the movement has been initiated and decisions about movement goals can be rapidly updated (Contemori et al., 2023; Nashed et al., 2014). We found that hand reaction times were shorter when intercepting targets at a distance of 3 compared to 6 *cm*. These results suggest that planning a shorter, or lower amplitude hand response took slightly less time (∼7 *ms*) than the longer, or higher amplitude response. This effect could be related to the observation that the hand responded more vigorously when intercepting targets at 6 compared to 3 *cm*, and thus higher muscle recruitment was required. Future work investigating muscle activity is required to elucidate this idea.

In contrast to hand responses, we found that eye reaction times were shorter for targets that jumped 6 *cm*, or 9.2 *deg*, compared to 3 *cm*, or 4.6 *deg*. This is in contrast to previous findings in human and non-human primates, showing that when making visually-guided saccades to targets that are further than 1 *deg* away, saccade latency increases with increasing target eccentricity (Hafed & Goffart, 2020; Kalesnykas & Hallett, 1994; Zhang & Fries, 2023). It should be noted that in these studies, saccades were made to stationary targets at ‘regular’ saccade latencies. Interestingly, short-latency saccades to predictable targets in the periphery are initiated earlier toward more eccentric compared to less eccentric targets (Cohen & Ross, 1977). Further, saccades to auditory targets are initiated earlier the more eccentric the target position (Gabriel et al., 2010). Taken together, these results suggest that salient sensory events may evoke even earlier orienting responses for targets of relatively high eccentricity.

## Conclusion

This paper highlights system-specific mechanisms guiding rapid eye and hand responses. We found that both eye and hand responses were similarly modulated by jump certainty, with reaction times decreasing when perturbations were highly predictable. However, we found that increasing levels of response urgency impacted eye and hand responses differently. Whereas hand responses scaled to the level of urgency, eye responses were relatively unaffected. We proposed a framework that links our behavioural results to the neural mechanisms involved in the visuomotor coordination of the eye and hand. This framework can potentially offer a foundation for future research on motor actions in the face of unpredictable changes in our dynamic world.

## Citation diversity statement

Recent work in several fields of science has identified a bias in citation practices such that papers from women and other minority scholars are under-cited relative to the number of such papers in the field (Bertolero et al., 2020; Caplar et al., 2017; Chatterjee & Werner, 2021; Dion et al., 2018; Dworkin et al., 2020; Fulvio et al., 2021; Maliniak et al., 2013; Mitchell et al., 2013; Wang et al., 2021). Here we sought to proactively consider choosing references that reflect the diversity of the field in thought, form of contribution, gender, race, ethnicity, and other factors. First, we obtained the predicted gender of the first and last author of each reference by using databases that store the probability of a first name being carried by a woman (Dworkin et al., 2020; Zhou et al., 2020). By this measure (and excluding self-citations to the first and last authors of our current paper), our references contain 7.14% woman(first)/woman(last), 8.16% man/woman, 19.39% woman/man, and 65.31% man/man. This method is limited in that a) names, pronouns, and social media profiles used to construct the databases may not, in every case, be indicative of gender identity and b) it cannot account for intersex, non-binary, or transgender people. Second, we obtained the predicted racial/ethnic category of the first and last author of each reference by databases that store the probability of a first and last name being carried by an author of colour (Ambekar et al., 2009; Chintalapati et al., 2023). By this measure (and excluding self-citations), our references contain 4.67% author of colour (first)/author of colour(last), 23.22% white author/author of colour, 17.89% author of colour/white author, and 54.22% white author/white author. This method is limited in that a) names and Florida Voter Data to make the predictions may not be indicative of racial/ethnic identity, and b) it cannot account for Indigenous and mixed-race authors, or those who may face differential biases due to the ambiguous racialization or ethnicization of their names. We look forward to future work that could help us to better understand how to support equitable practices in science.

## Credit authorship contribution statement

Conceptualization (JF, PB, KP, JRF, SHS), Methodology (JF, PB, KP, SHS), Software (PB, KP), Validation (PB, KP, SHS), Formal analysis (KP, JF, SHS), Resources (JRF, SHS), Data Curation (JF), Writing - Original Draft (JF, PB, KP), Writing - review & editing (JF, PB, KP, JRF, SHS), Visualization (JF), Supervision (JF, JRF, SHS), Funding acquisition (JF, JRF, SHS)

## Acknowledgements

This work was supported by by a Deutsche Forschungsgemeinschaft (DFG) Research Fellowship to JF (grant FO 1347/1-1), an NSERC Discovery grant to JRF, and an NSERC Discovery grant to SHS. The authors thank Elissa Robichaud and Nethmi Illamperuma for their help with data collection.

## References

Ambekar, A., Ward, C., Mohammed, J., Male, S., & Skiena, S. (2009). Name-ethnicity classification from open sources. Proceedings of the 15th ACM SIGKDD International Conference on Knowledge Discovery and Data Mining, 49–58. 10.1145/1557019.1557032

Barany, D. A., Gómez-Granados, A., Schrayer, M., Cutts, S. A., & Singh, T. (2020). Perceptual decisions about object shape bias visuomotor coordination during rapid interception movements. Journal of Neurophysiology, 123(6), 2235–2248. 10.1152/jn.00098.2020

Battaglia-Mayer, A., Caminiti, R., Lacquaniti, F., & Zago, M. (2003). Multiple Levels of Representation of Reaching in the Parieto-frontal Network. Cerebral Cortex, 13(10), 1009–1022. 10.1093/cercor/13.10.1009

Bertolero, M. A., Dworkin, J. D., David, S. U., Lloreda, C. L., Srivastava, P., Stiso, J., Zhou, D., Dzirasa, K., Fair, D. A., Kaczkurkin, A. N., Marlin, B. J., Shohamy, D., Uddin, L. Q., Zurn, P., & Bassett, D. S. (2020). Racial and ethnic imbalance in neuroscience reference lists and intersections with gender. 10.1101/2020.10.12.336230

Binaee, K., & Diaz, G. (2019). Movements of the eyes and hands are coordinated by a common predictive strategy. Journal of Vision, 19(12), 3. 10.1167/19.12.3

Boehnke, S. E., & Munoz, D. P. (2008). On the importance of the transient visual response in the superior colliculus. Current Opinion in Neurobiology, 18(6), 544–551. 10.1016/j.conb.2008.11.004

Bosco, G., Delle Monache, S., Gravano, S., Indovina, I., La Scaleia, B., Maffei, V., Zago, M., & Lacquaniti, F. (2015). Filling gaps in visual motion for target capture. Frontiers in Integrative Neuroscience, 9. 10.3389/fnint.2015.00013

Brenner, E., & Smeets, J. B. J. (1997). Fast Responses of the Human Hand to Changes in Target Position. Journal of Motor Behavior, 29(4), 297–310. 10.1080/00222899709600017

Brenner, E., & Smeets, J. B. J. (2018). Continuously updating one’s predictions underlies successful interception. Journal of Neurophysiology, 120(6), 3257–3274. 10.1152/jn.00517.2018

Caplar, N., Tacchella, S., & Birrer, S. (2017). Quantitative evaluation of gender bias in astronomical publications from citation counts. Nature Astronomy, 1(6), 0141. 10.1038/s41550-017-0141

Carpenter, R. H. S., & Williams, M. L. L. (1995). Neural computation of log likelihood in control of saccadic eye movements. Nature, 377(6544), 59–62. 10.1038/377059a0

Cesqui, B., Mezzetti, M., Lacquaniti, F., & d’Avella, A. (2015). Gaze Behavior in One-Handed Catching and Its Relation with Interceptive Performance: What the Eyes Can’t Tell. PLOS ONE, 10(3), e0119445. 10.1371/journal.pone.0119445

Chatterjee, P., & Werner, R. M. (2021). Gender Disparity in Citations in High-Impact Journal Articles. JAMA Network Open, 4(7), e2114509. 10.1001/jamanetworkopen.2021.14509

Chintalapati, R., Laohaprapanon, S., & Sood, G. (2023). Predicting Race and Ethnicity From the Sequence of Characters in a Name (No. arXiv:1805.02109). arXiv. http://arxiv.org/abs/1805.02109

Coe, B. C., & Munoz, D. P. (2017). Mechanisms of saccade suppression revealed in the anti-saccade task. Philosophical Transactions of the Royal Society B: Biological Sciences, 372(1718), 20160192. 10.1098/rstb.2016.0192

Cohen, M. E., & Ross, L. E. (1977). Saccade latency in children and adults: Effects of warning interval and target eccentricity. Journal of Experimental Child Psychology, 23(3), 539–549. 10.1016/0022-0965(77)90044-3

Contemori, S., Loeb, G. E., Corneil, B. D., Wallis, G., & Carroll, T. J. (2021). The influence of temporal predictability on express visuomotor responses. Journal of Neurophysiology, 125(3), 731–747. 10.1152/jn.00521.2020

Contemori, S., Loeb, G. E., Corneil, B. D., Wallis, G., & Carroll, T. J. (2022). Symbolic cues enhance express visuomotor responses in human arm muscles at the motor planning rather than the visuospatial processing stage. Journal of Neurophysiology, 128(3), 494–510. 10.1152/jn.00136.2022

Contemori, S., Loeb, G. E., Corneil, B. D., Wallis, G., & Carroll, T. J. (2023). Express Visuomotor Responses Reflect Knowledge of Both Target Locations and Contextual Rules during Reaches of Different Amplitudes. The Journal of Neuroscience, 43(42), 7041–7055. 10.1523/JNEUROSCI.2069-22.2023

Cooper, B., & McPeek, R. M. (2021). Role of the Superior Colliculus in Guiding Movements Not Made by the Eyes. Annual Review of Vision Science, 7(1), 279–300. 10.1146/annurev-vision-012521-102314

Corneil, B. D., & Munoz, D. P. (2014). Overt Responses during Covert Orienting. Neuron, 82(6), 1230–1243. 10.1016/j.neuron.2014.05.040

Corneil, B. D., Munoz, D. P., Chapman, B. B., Admans, T., & Cushing, S. L. (2008). Neuromuscular consequences of reflexive covert orienting. Nature Neuroscience, 11(1), 13–15. 10.1038/nn2023

Corneil, B. D., Olivier, E., & Munoz, D. P. (2004). Visual Responses on Neck Muscles Reveal Selective Gating that Prevents Express Saccades. Neuron, 42(5), 831–841. 10.1016/S0896-6273(04)00267-3

Coudiere, A., & Danion, F. R. (2024). Eye-hand coordination all the way: From discrete to continuous hand movements. Journal of Neurophysiology, 131(4), 652–667. 10.1152/jn.00314.2023

Crawford, J. D., Medendorp, W. P., & Marotta, J. J. (2004). Spatial Transformations for Eye– Hand Coordination. Journal of Neurophysiology, 92(1), 10–19. 10.1152/jn.00117.2004

Crévécoeur, F., Kurtzer, I., Bourke, T., & Scott, S. H. (2013). Feedback responses rapidly scale with the urgency to correct for external perturbations. Journal of Neurophysiology, 110(6), 1323–1332. 10.1152/jn.00216.2013

Cross, K. P., Cluff, T., Takei, T., & Scott, S. H. (2019). Visual Feedback Processing of the Limb Involves Two Distinct Phases. The Journal of Neuroscience, 39(34), 6751–6765. 10.1523/JNEUROSCI.3112-18.2019

de Brouwer, A. J., Flanagan, J. R., & Spering, M. (2021). Functional Use of Eye Movements for an Acting System. Trends in Cognitive Sciences, 25(3), 252–263. 10.1016/j.tics.2020.12.006

de la Malla, C., Rushton, S. K., Clark, K., Smeets, J. B. J., & Brenner, E. (2019). The predictability of a target’s motion influences gaze, head, and hand movements when trying to intercept it. Journal of Neurophysiology, 121(6), 2416–2427. 10.1152/jn.00917.2017

Dean, H. L., Hagan, M. A., & Pesaran, B. (2012). Only Coherent Spiking in Posterior Parietal Cortex Coordinates Looking and Reaching. Neuron, 73(4), 829–841. 10.1016/j.neuron.2011.12.035

Diaz, G. J., Cooper, J., Rothkopf, C., & Hayhoe, M. M. (2013). Saccades to future ball location reveal memory-based prediction in a virtual-reality interception task. Journal of Vision, 13(1), 20–20. 10.1167/13.1.20

Dimitriou, M., Wolpert, D. M., & Franklin, D. W. (2013). The Temporal Evolution of Feedback Gains Rapidly Update to Task Demands. Journal of Neuroscience, 33(26), 10898–10909. 10.1523/JNEUROSCI.5669-12.2013

Dion, M. L., Sumner, J. L., & Mitchell, S. M. (2018). Gendered Citation Patterns across Political Science and Social Science Methodology Fields. Political Analysis, 26(3), 312–327. 10.1017/pan.2018.12

Dorris, M. C., Klein, R. M., Everling, S., & Munoz, D. P. (2002). Contribution of the Primate Superior Colliculus to Inhibition of Return. Journal of Cognitive Neuroscience, 14(8), 1256–1263. 10.1162/089892902760807249

Dorris, M. C., Paré, M., & Munoz, D. P. (1997). Neuronal Activity in Monkey Superior Colliculus Related to the Initiation of Saccadic Eye Movements. The Journal of Neuroscience, 17(21), 8566–8579. 10.1523/JNEUROSCI.17-21-08566.1997

Dworkin, J. D., Linn, K. A., Teich, E. G., Zurn, P., Shinohara, R. T., & Bassett, D. S. (2020). The extent and drivers of gender imbalance in neuroscience reference lists. Nature Neuroscience, 23(8), 918–926. 10.1038/s41593-020-0658-y

Engel, K. C., Anderson, J. H., & Soechting, J. F. (1999). Oculomotor Tracking in Two Dimensions. Journal of Neurophysiology, 81(4), 1597–1602. 10.1152/jn.1999.81.4.1597

Etchells, P. J., Benton, C. P., Ludwig, C. J. H., & Gilchrist, I. D. (2010). The target velocity integration function for saccades. Journal of Vision, 10(6), 7–7. 10.1167/10.6.7

Findlay, J. M. (1983). Visual Information Processing for Saccadic Eye Movements. In A. Hein & M. Jeannerod (Eds.), Spatially Oriented Behavior (pp. 281–303). Springer New York. 10.1007/978-1-4612-5488-1_16

Fischer, B., & Ramsperger, E. (1984). Human express saccades: Extremely short reaction times of goal directed eye movements. Experimental Brain Research, 57(1). 10.1007/BF00231145

Fisk, J. D., & Goodale, M. A. (1985). The organization of eye and limb movements during unrestricted reaching to targets in contralateral and ipsilateral visual space. Experimental Brain Research, 60(1). 10.1007/BF00237028

Fooken, J., Baltaretu, B. R., Barany, D. A., Diaz, G., Semrau, J. A., Singh, T., & Crawford, J. D. (2023). Perceptual-Cognitive Integration for Goal-Directed Action in Naturalistic Environments. Journal of Neuroscience, 43(45), 7511–7522. 10.1523/JNEUROSCI.1373-23.2023

Fooken, J., Kreyenmeier, P., & Spering, M. (2021). The role of eye movements in manual interception: A mini-review. Vision Research, 183, 81–90. 10.1016/j.visres.2021.02.007

Fooken, J., & Spering, M. (2019). Decoding go/no-go decisions from eye movements. Journal of Vision, 19(2), 5. 10.1167/19.2.5

Fooken, J., & Spering, M. (2020). Eye movements as a readout of sensorimotor decision processes. Journal of Neurophysiology, 123(4), 1439–1447. 10.1152/jn.00622.2019

Fooken, J., Yeo, S.-H., Pai, D. K., & Spering, M. (2016). Eye movement accuracy determines natural interception strategies. Journal of Vision, 16(14), 1. 10.1167/16.14.1

Frens, M., & Erkelens, C. J. (1991). Coordination of hand movements and saccades: Evidence for a common and a separate pathway. Experimental Brain Research, 85(3). 10.1007/BF00231754

Fulvio, J. M., Akinnola, I., & Postle, B. R. (2021). Gender (Im)balance in Citation Practices in Cognitive Neuroscience. Journal of Cognitive Neuroscience, 33(1), 3–7. 10.1162/jocn_a_01643

Gabriel, D. N., Munoz, D. P., & Boehnke, S. E. (2010). The eccentricity effect for auditory saccadic reaction times is independent of target frequency. Hearing Research, 262(1–2), 19–25. 10.1016/j.heares.2010.01.016

Gandhi, N. J., & Katnani, H. A. (2011). Motor Functions of the Superior Colliculus. Annual Review of Neuroscience, 34(1), 205–231. 10.1146/annurev-neuro-061010-113728

Gellman, R. S., & Carl, J. R. (1991). Motion processing for saccadic eye movements in humans. Experimental Brain Research, 84(3). 10.1007/BF00230979

Goettker, A., Braun, D. I., & Gegenfurtner, K. R. (2019). Dynamic combination of position and motion information when tracking moving targets. Journal of Vision, 19(7), 2. 10.1167/19.7.2

Goettker, A., Brenner, E., Gegenfurtner, K. R., & de la Malla, C. (2019). Corrective saccades influence velocity judgments and interception. Scientific Reports, 9(1), 5395. 10.1038/s41598-019-41857-z

Goonetilleke, S. C., Katz, L., Wood, D. K., Gu, C., Huk, A. C., & Corneil, B. D. (2015). Cross-species comparison of anticipatory and stimulus-driven neck muscle activity well before saccadic gaze shifts in humans and nonhuman primates. Journal of Neurophysiology, 114(2), 902–913. 10.1152/jn.00230.2015

Gribble, P. L., Everling, S., Ford, K., & Mattar, A. (2002). Hand-eye coordination for rapid pointing movements: Arm movement direction and distance are specified prior to saccade onset. Experimental Brain Research, 145(3), 372–382. 10.1007/s00221-002-1122-9

Gu, C., Wood, D. K., Gribble, P. L., & Corneil, B. D. (2016). A Trial-by-Trial Window into Sensorimotor Transformations in the Human Motor Periphery. Journal of Neuroscience, 36(31), 8273–8282. 10.1523/JNEUROSCI.0899-16.2016

Hafed, Z. M., & Goffart, L. (2020). Gaze direction as equilibrium: More evidence from spatial and temporal aspects of small-saccade triggering in the rhesus macaque monkey. Journal of Neurophysiology, 123(1), 308–322. 10.1152/jn.00588.2019

Hwang, E. J., Hauschild, M., Wilke, M., & Andersen, R. A. (2014). Spatial and Temporal Eye-Hand Coordination Relies on the Parietal Reach Region. Journal of Neuroscience, 34(38), 12884–12892. 10.1523/JNEUROSCI.3719-13.2014

Irving, E. L., Steinbach, M. J., Lillakas, L., Babu, R. J., & Hutchings, N. (2006). Horizontal Saccade Dynamics across the Human Life Span. Investigative Opthalmology & Visual Science, 47(6), 2478. 10.1167/iovs.05-1311

Jakobs, O., Wang, L. E., Dafotakis, M., Grefkes, C., Zilles, K., & Eickhoff, S. B. (2009). Effects of timing and movement uncertainty implicate the temporo-parietal junction in the prediction of forthcoming motor actions. NeuroImage, 47(2), 667–677. 10.1016/j.neuroimage.2009.04.065

Johansson, R. S., Westling, G., Bäckström, A., & Flanagan, J. R. (2001). Eye–Hand Coordination in Object Manipulation. The Journal of Neuroscience, 21(17), 6917–6932. 10.1523/JNEUROSCI.21-17-06917.2001

Kalesnykas, R. P., & Hallett, P. E. (1994). Retinal eccentricity and the latency of eye saccades. Vision Research, 34(4), 517–531. 10.1016/0042-6989(94)90165-1

Kalidindi, H. T., & Crévécoeur, F. (2023). Human reaching control in dynamic environments. Current Opinion in Neurobiology, 83, 102810. 10.1016/j.conb.2023.102810

Kang, J. U., Mooshagian, E., & Snyder, L. H. (2024). Functional organization of posterior parietal cortex circuitry based on inferred information flow. Cell Reports, 43(4), 114028. 10.1016/j.celrep.2024.114028

Kingstone, A., & Klein, R. M. (1993). What are human express saccades? Perception & Psychophysics, 54(2), 260–273. 10.3758/BF03211762

Korbisch, C. C., Apuan, D. R., Shadmehr, R., & Ahmed, A. A. (2022). Saccade vigor reflects the rise of decision variables during deliberation. Current Biology, 32(24), 5374–5381.e4. 10.1016/j.cub.2022.10.053

Kozak, R. A., Cecala, A. L., & Corneil, B. D. (2020). An Emerging Target Paradigm to Evoke Fast Visuomotor Responses on Human Upper Limb Muscles. Journal of Visualized Experiments, 162, 61428. 10.3791/61428

Kozak, R. A., Kreyenmeier, P., Gu, C., Johnston, K., & Corneil, B. D. (2019). Stimulus-Locked Responses on Human Upper Limb Muscles and Corrective Reaches Are Preferentially Evoked by Low Spatial Frequencies. eNeuro, 6(5), ENEURO.0301-19.2019. 10.1523/ENEURO.0301-19.2019

Kreyenmeier, P., Kämmer, L., Fooken, J., & Spering, M. (2022). Humans Can Track But Fail to Predict Accelerating Objects. Eneuro, 9(5), ENEURO.0185-22.2022. 10.1523/ENEURO.0185-22.2022

Land, M. F. (2006). Eye movements and the control of actions in everyday life. Progress in Retinal and Eye Research, 25(3), 296–324. 10.1016/j.preteyeres.2006.01.002

Land, M. F., & McLeod, P. (2000). From eye movements to actions: How batsmen hit the ball. Nature Neuroscience, 3(12), 1340–1345. 10.1038/81887

Maliniak, D., Powers, R., & Walter, B. F. (2013). The Gender Citation Gap in International Relations. International Organization, 67(4), 889–922. 10.1017/S0020818313000209

Mann, D. L., Nakamoto, H., Logt, N., Sikkink, L., & Brenner, E. (2019). Predictive eye movements when hitting a bouncing ball. Journal of Vision, 19(14), 28. 10.1167/19.14.28

Marino, R. A., Levy, R., Boehnke, S., White, B. J., Itti, L., & Munoz, D. P. (2012). Linking visual response properties in the superior colliculus to saccade behavior. European Journal of Neuroscience, 35(11), 1738–1752. 10.1111/j.1460-9568.2012.08079.x

Marino, R. A., Levy, R., & Munoz, D. P. (2015). Linking express saccade occurance to stimulus properties and sensorimotor integration in the superior colliculus. Journal of Neurophysiology, 114(2), 879–892. 10.1152/jn.00047.2015

Maurus, P., Jackson, K., Cashaback, J. G. A., & Cluff, T. (2023). The nervous system tunes sensorimotor gains when reaching in variable mechanical environments. iScience, 26(6), 106756. 10.1016/j.isci.2023.106756

Mitchell, S. M., Lange, S., & Brus, H. (2013). Gendered Citation Patterns in International Relations Journals. International Studies Perspectives, 14(4), 485–492. 10.1111/insp.12026

Mrotek, L. A. (2013). Following and intercepting scribbles: Interactions between eye and hand control. Experimental Brain Research, 227(2), 161–174. 10.1007/s00221-013-3496-2

Mrotek, L. A., & Soechting, J. F. (2007). Target Interception: Hand–Eye Coordination and Strategies. The Journal of Neuroscience, 27(27), 7297–7309. 10.1523/JNEUROSCI.2046-07.2007

Nashed, J. Y., Crévécoeur, F., & Scott, S. H. (2014). Rapid Online Selection between Multiple Motor Plans. The Journal of Neuroscience, 34(5), 1769–1780. 10.1523/JNEUROSCI.3063-13.2014

Paré, M., & Munoz, D. P. (1996). Saccadic reaction time in the monkey: Advanced preparation of oculomotor programs is primarily responsible for express saccade occurrence. Journal of Neurophysiology, 76(6), 3666–3681. 10.1152/jn.1996.76.6.3666

Park, K., Ritsma, B. R., Dukelow, S. P., & Scott, S. H. (2023). A robot-based interception task to quantify upper limb impairments in proprioceptive and visual feedback after stroke. Journal of NeuroEngineering and Rehabilitation, 20(1), 137. 10.1186/s12984-023-01262-0

Perfiliev, S., Isa, T., Johnels, B., Steg, G., & Wessberg, J. (2010). Reflexive Limb Selection and Control of Reach Direction to Moving Targets in Cats, Monkeys, and Humans. Journal of Neurophysiology, 104(5), 2423–2432. 10.1152/jn.01133.2009

Philipp, R., & Hoffmann, K.-P. (2014). Arm Movements Induced by Electrical Microstimulation in the Superior Colliculus of the Macaque Monkey. The Journal of Neuroscience, 34(9), 3350–3363. 10.1523/JNEUROSCI.0443-13.2014

Poscente, S. V., Peters, R. M., Cashaback, J. G. A., & Cluff, T. (2021). Rapid Feedback Responses Parallel the Urgency of Voluntary Reaching Movements. Neuroscience, 475, 163–184. 10.1016/j.neuroscience.2021.07.014

Prablanc, C., Echallier, J. F., Komilis, E., & Jeannerod, M. (1979). Optimal response of eye and hand motor systems in pointing at a visual target: I. Spatio-temporal characteristics of eye and hand movements and their relationships when varying the amount of visual information. Biological Cybernetics, 35(2), 113–124. 10.1007/BF00337436

Pruszynski, J. A., King, G. L., Boisse, L., Scott, S. H., Flanagan, J. R., & Munoz, D. P. (2010). Stimulus-locked responses on human arm muscles reveal a rapid neural pathway linking visual input to arm motor output. European Journal of Neuroscience, 32(6), 1049–1057. 10.1111/j.1460-9568.2010.07380.x

Reppert, T. R., Lempert, K. M., Glimcher, P. W., & Shadmehr, R. (2015). Modulation of Saccade Vigor during Value-Based Decision Making. The Journal of Neuroscience, 35(46), 15369–15378. 10.1523/JNEUROSCI.2621-15.2015

Reschechtko, S., Gnanaseelan, C., & Pruszynski, J. A. (2023). Reach Corrections Toward Moving Objects are Faster Than Reach Corrections Toward Instantaneously Switching Targets. Neuroscience, 526, 135–143. 10.1016/j.neuroscience.2023.06.021

Robinson, D. A. (2022). Behavior of the saccadic system: Metrics of timing and accuracy. In Progress in Brain Research (Vol. 267, pp. 329–353). Elsevier. 10.1016/bs.pbr.2021.10.016

Ron, S., Vieville, T., & Droulez, J. (1989). Target velocity based prediction in saccadic vector programming. Vision Research, 29(9), 1103–1114. 10.1016/0042-6989(89)90059-X

Sailer, U., Eggert, T., Ditterich, J., & Straube, A. (2000). Spatial and temporal aspects of eye-hand coordination across different tasks. Experimental Brain Research, 134(2), 163–173. 10.1007/s002210000457

Salinas, E., Steinberg, B. R., Sussman, L. A., Fry, S. M., Hauser, C. K., Anderson, D. D., & Stanford, T. R. (2019). Voluntary and involuntary contributions to perceptually guided saccadic choices resolved with millisecond precision. eLife, 8, e46359. 10.7554/eLife.46359

Saslow, M. G. (1967). Effects of Components of Displacement-Step Stimuli Upon Latency for Saccadic Eye Movement. Journal of the Optical Society of America, 57(8), 1024. 10.1364/JOSA.57.001024

Schlag, J., & Schlag-Rey, M. (2002). Through the eye, slowly; Delays and localization errors in the visual system. Nature Reviews Neuroscience, 3(3), 191–191. 10.1038/nrn750

Scott, S. H. (2016). A Functional Taxonomy of Bottom-Up Sensory Feedback Processing for Motor Actions. Trends in Neurosciences, 39(8), 512–526. 10.1016/j.tins.2016.06.001

Seideman, J. A., Stanford, T. R., & Salinas, E. (2018). Saccade metrics reflect decision-making dynamics during urgent choices. Nature Communications, 9(1), 2907. 10.1038/s41467-018-05319-w

Shadmehr, R., Reppert, T. R., Summerside, E. M., Yoon, T., & Ahmed, A. A. (2019). Movement Vigor as a Reflection of Subjective Economic Utility. Trends in Neurosciences, 42(5), 323–336. 10.1016/j.tins.2019.02.003

Sparks, D., Rohrer, W. H., & Zhang, Y. (2000). The role of the superior colliculus in saccade initiation: A study of express saccades and the gap effect. Vision Research, 40(20), 2763–2777. 10.1016/S0042-6989(00)00133-4

Spering, M., Schütz, A. C., Braun, D. I., & Gegenfurtner, K. R. (2011). Keep your eyes on the ball: Smooth pursuit eye movements enhance prediction of visual motion. Journal of Neurophysiology, 105(4), 1756–1767. 10.1152/jn.00344.2010

Stanford, T. R., & Salinas, E. (2021). Urgent Decision Making: Resolving Visuomotor Interactions at High Temporal Resolution. Annual Review of Vision Science, 7(1), 323–348. 10.1146/annurev-vision-100419-103842

Stanford, T. R., Shankar, S., Massoglia, D. P., Costello, M. G., & Salinas, E. (2010). Perceptual decision making in less than 30 milliseconds. Nature Neuroscience, 13(3), 379–385. 10.1038/nn.2485

Stuphorn, V., Hoffmann, K.-P., & Miller, L. E. (1999). Correlation of Primate Superior Colliculus and Reticular Formation Discharge With Proximal Limb Muscle Activity. Journal of Neurophysiology, 81(4), 1978–1982. 10.1152/jn.1999.81.4.1978

Todorov, E., & Jordan, M. I. (2002). Optimal feedback control as a theory of motor coordination. Nature Neuroscience, 5(11), 1226–1235. 10.1038/nn963

Veerman, M. M., Brenner, E., & Smeets, J. B. J. (2008). The latency for correcting a movement depends on the visual attribute that defines the target. Experimental Brain Research, 187(2), 219–228. 10.1007/s00221-008-1296-x

Vesia, M., & Crawford, J. D. (2012). Specialization of reach function in human posterior parietal cortex. Experimental Brain Research, 221(1), 1–18. 10.1007/s00221-012-3158-9

Wang, X., Dworkin, J. D., Zhou, D., Stiso, J., Falk, E. B., Bassett, D. S., Zurn, P., & Lydon-Staley, D. M. (2021). Gendered citation practices in the field of communication. Annals of the International Communication Association, 45(2), 134–153. 10.1080/23808985.2021.1960180

Werner, W., Dannenberg, S., & Hoffmann, K.-P. (1997). Arm-movement-related neurons in the primate superior colliculus and underlying reticular formation: Comparison of neuronal activity with EMGs of muscles of the shoulder, arm and trunk during reaching: Experimental Brain Research, 115(2), 191–205. 10.1007/PL00005690

Wood, D. K., Gu, C., Corneil, B. D., Gribble, P. L., & Goodale, M. A. (2015). Transient visual responses reset the phase of low-frequency oscillations in the skeletomotor periphery. European Journal of Neuroscience, 42(3), 1919–1932. 10.1111/ejn.12976

Yoon, T., Jaleel, A., Ahmed, A. A., & Shadmehr, R. (2020). Saccade vigor and the subjective economic value of visual stimuli. Journal of Neurophysiology, 123(6), 2161–2172. 10.1152/jn.00700.2019

Zago, M., Iosa, M., Maffei, V., & Lacquaniti, F. (2010). Extrapolation of vertical target motion through a brief visual occlusion. Experimental Brain Research, 201(3), 365–384. 10.1007/s00221-009-2041-9

Zago, M., McIntyre, J., Senot, P., & Lacquaniti, F. (2009). Visuo-motor coordination and internal models for object interception. Experimental Brain Research, 192, 571–604.

Zhang, Y., & Fries, P. (2023). Eccentricity-Dependent Saccadic Reaction Time: The Roles of Foveal Magnification and Attentional Orienting. 10.1101/2023.08.08.552339

Zhou, D., Bertolero, M. A., Stiso, J., Cornblath, E. J., Teich, E. G., Blevins, A. S., Virtualmario Camp, Dworkin, J., & Bassett, D. S. (2020). Gender diversity statement and code notebook (v1.1) [Computer software]. https://github.com/dalejn/cleanBib

